# Integrating roots into a whole plant network of flowering time genes in *Arabidopsis thaliana*

**DOI:** 10.1101/036244

**Authors:** Frédéric Bouché, Maria D’Aloia, Pierre Tocquin, Guillaume Lobet, Nathalie Detry, Claire Périlleux

## Abstract

Molecular data concerning the involvement of the roots in the genetic pathways regulating floral transition are lacking. In this study, we performed global analyses of root transcriptome in Arabidopsis in order to identify flowering time genes that are expressed in the roots and genes that are differentially expressed in the roots during the induction of flowering. Data mining of public microarray experiments uncovered that about 200 genes whose mutation was reported to alter flowering time are expressed in the roots but only few flowering integrators were found. Transcriptomic analysis of the roots during synchronized induction of flowering by a single 22-h long day revealed that 595 genes were differentially expressed. A delay in clock gene expression was observed upon extension of the photoperiod. Enrichment analyses of differentially expressed genes in root tissues, gene ontology categories and cis-regulatory elements converged towards sugar signaling. We inferred that roots are integrated in systemic signaling whereby carbon supply coordinates growth at the whole plant level during the induction of flowering.

## INTRODUCTION

Flowering is a crucial step of plant development that must be precisely timed to occur when external conditions are favourable for successful reproduction. Floral induction is therefore controlled by several environmental and endogenous cues, whose inputs are integrated into finely-tuned regulatory gene networks. In *Arabidopsis thaliana,* genetic analyses unveiled a number of flowering pathways that are activated in response to photoperiod, temperature, sugars, hormones and plant aging, or eventually occurs autonomously (Bouché et al., 2016). These pathways are not restricted to the shoot apical meristem where flowers are initiated but also involve the leaves at least, supporting the fact that flowering, as shown previously at the physiological level, is a systemic process. Clearest evidence came from the photoperiodic pathway that accelerates flowering in response to increasing daylength to ensure bolting in spring (Song et al., 2015). A key actor in this pathway is the transcription factor CONSTANS (CO) whose expression follows a circadian pattern but is degraded in the dark (Suarez-Lopez et al., 2001; Valverde et al., 2004). Light must therefore coincide with CO synthesis to stabilize the protein and enable activation of its targets (Valverde et al., 2004). This occurs during long days in the companion cells of phloem, where CO activates *FLOWERING LOCUS T (FT)* (Samach et al., 2000). The FT protein then moves systemically (Corbesier et al., 2007) and in the shoot apical meristem interacts with the transcription factor FD via 14-3-3 proteins (Abe et al., 2005; Wigge, 2005; Taoka et al., 2011). This flowering activation complex triggers the expression of genes that are responsible for the conversion of the vegetative shoot apical meristem into an inflorescence meristem and for the promotion of floral fate in lateral primordia (Ó’Maoiléidigh et al., 2014).

The prominent role of the FT protein in the systemic signaling operating at floral transition opens questions concerning the role of side molecules that are co-transported from leaf sources in the phloem and the pleiotropic effects of FT and putative co-transported signals in different sinks. Sugar loading is the first step of long-distance mass-flow movement in phloem and hence carbohydrates might influence flowering signals delivery (Dinant and Suarez-Lopez, 2012). Several reports however indicate that sugars act as flowering signals themselves, at two sites in the plant. In the leaves, photosynthesis and activity of TREHALOSE-6-PHOSPHATE SYNTHASE 1 (TPS1), which catalyzes the formation of trehalose-6-phosphate (T6P) involved in sugar sensing, are required for the induction of the *FT* gene, even under inductive photoperiod (King et al., 2008; Wahl et al., 2013). The plant so integrates an environmental signal (the activation of *FT* by CO in response to increasing day length) with a physiological signal (the presence of high carbohydrate levels, as indicated by T6P) (Wahl et al., 2013). Interestingly, CO regulates the expression of GRANULE-BOUND STARCH SYNTHASE (GBSS), an enzyme controlling the synthesis of amylose in starch granules, and could thereby mediate modification of transitory starch composition to increase the sugar mobilization at floral transition (Ortiz-Marchena et al., 2014). Using starchless mutant, Corbesier et al. (1998) concluded that starch mobilization was critical for floral induction in conditions which did not involve an increased photosynthetic activity. All those results build evidence for sugar contribution to the florigenic signaling. In the shoot apex, sucrose content increases when Arabidopsis plant flowers in response to a photosynthetic long day (King et al., 2008; Corbesier et al., 1998) or eventually in short days (Eriksson et al., 2006). Sugars can induce the expression of flowering genes in the meristem, *e.g.* via the T6P pathway, independently of *FT* (Wahl et al., 2013). Beside sugars, the phloem sap of Arabidopsis is also enriched in amino acids and hormones of the cytokinin family when flowering is induced by a photoperiodic change (Corbesier et al., 1998; 2003). Cytokinins can promote flowering by inducing the paralogue of *FT, TWIN SISTER OF FT (TSF),* in the leaves and downstream flowering genes in the shoot apical meristem (D’Aloia et al., 2011).

If we can infer from the previous section that multiple flowering signals including FT, sugars and hormones are transported in phloem, the signaling route appears as simplified to one way from leaves to the shoot apical meristem. Roots are ignored. At the physiological level though, a shoot-to-root-to-shoot loop has been described to drive sugar and cytokinin fluxes at floral transition in the Arabidopsis relative white mustard (Havelange et al., 2000). More directly, tagging of the FT protein with GFP in Arabidopsis allowed to detect movement of the fusion protein from overexpressor scion to *ft* mutant rootstock, indicating that it is not restricted to aerial parts of the plant (Corbesier et al., 2007). In other species, FT-like proteins exported from the leaves can induce belowground processes such as tuberization in potato (Navarro et al., 2011) or bulb formation in onion (Lee et al., 2013). These reports indicate that developmental signals from leaf origin reach the underground organs.

Little information is available about the expression of flowering time genes in the roots. In a few cases only, analysis of expression patterns or phenotyping of mutants included careful examination of the roots and were followed by complementation tests (Bernier and Périlleux, 2005). This rationale was used for *FT,* which is not expressed in the roots but whose partner *FD* is (Abe et al., 2005), raising the possibility of a role for the flowering activation complex in the roots. However, the root-specific expression of *FT* did not rescue the phenotype of *ft* single mutant, indicating that the expression of *FT* in root tissues is not sufficient – albeit it might contribute – to flowering (Abe et al., 2005). Other flowering time mutants such as *fca,* several *squamosa-promoter binding protein like (spl3, spl9* and *spl10)* or *terminal flower 1 (tfl1)* show root architecture phenotypes (Macknight et al., 2002; Lachowiec et al., 2015; Yu et al., 2015). Major flowering QTL in *Arabidopsis* were also found to be associated with root xylem secondary growth (Sibout et al., 2008). However, whether those traits indicate root-specific functions or indirect effects of flowering time genes remains to be demonstrated.

The aim of this study was to clarify the role of roots in the flowering process. Two complementary approaches were used. First, data mining of public microarray databases was performed to obtain a global view of flowering-time genes expressed in the roots. Second, the transcriptome of the roots was analysed during the induction of flowering. The set of differential by expressed genes was crossed with publicly available datasets obtained in different contexts for discovering potential regulatory networks.

## RESULTS

### A majority of flowering-time genes are expressed in roots

Data mining was performed using transcriptomic analyses of roots that are available in the ArrayExpress repository (Kolesnikov et al., 2015) (Figure 1). The whole set of selected experiments contained 1,673 Arabidopsis ATH1 Genome arrays (Supplemental Table 1). For each array, we performed an Affymetrix present/absent call to identify root-expressed genes. Genes were considered as being expressed when transcripts were detected (p<0.01) in at least 50% of the 1,673 arrays. We crossed the results of this filter with a comprehensive list of 306 flowering-time genes that we established in the FLOR-ID database (Bouché et al., 2016). These genes are allocated among different pathways whereby flowering occurs in response to photoperiod, vernalization, aging, ambient temperature, hormones or sugar; an “autonomous pathway” leads to flowering independently of these signals and involves regulators of general processes such as chromatin remodeling, transcriptional machinery or proteasome activity. Eight genes under the control of several converging pathways are defined as “flowering-time integrators”: FT, *TSF, SUPPRESSOR OF OVEREXPRESSION OF COI (SOCI), AGAMOUS-LIKE 24 (AGL24), FRUITFULL (FUL), FLOWERING LOCUS C (FLC), SHORT VEGETATIVE PHASE (SVP)* and *LEAFY (LFY).* Given the design of ATH1 microarrays, 37 flowering-time genes including 11 genes encoding microRNAs could not be included in our survey because they are not represented in the probe set. Out of the 269 represented flowering time genes, 183 (68%) were expressed in roots in more than half of the analyzed arrays (Figure 1A; Supplemental Table 2). Some flowering pathways were more enriched than others (Figure 1B), *e.g.* the photoperiodic pathway with 70% of its genes being expressed in the roots or the sugar pathway with 7 genes out of 9 being active in the roots, including *TPS1.* As expected, genes controlling autonomous flowering via general regulatory processes were widely detected in roots (80%). A side category of circadian clock genes was also highlighted in the analysis. By contrast, a low proportion of genes from the hormones and aging pathways could be detected.

**Figure 1.**
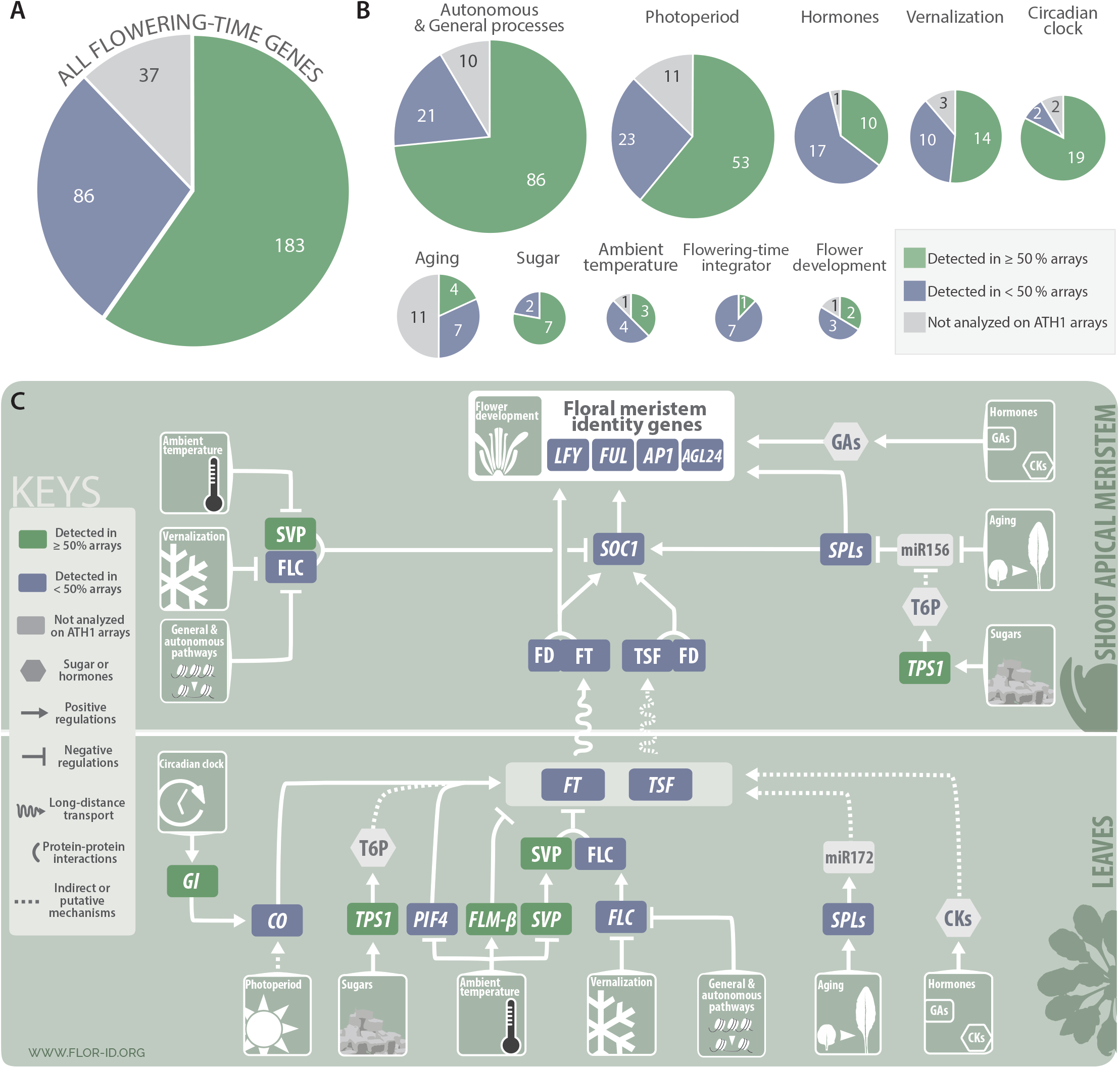
Flowering-time genes expressed in the roots of Arabidopsis thaliana. Root-expressed genes were identified by a present/absent call on 1,673 root ATH1 arrays retrieved from ArrayExpress repository (https://www.ebi.ac.uk/arrayexpress/). Flowering-time genes were extracted from FLOR-ID. **A**. All 306 flowering genes. **B**. Pie charts showing the same set of genes classified into flowering time pathways, circadian clock and flower development. Some genes are involved in more than one pathway. Pie chart area is proportional to gene number. **C**. The snapshot of flowering pathways was extracted and adapted from FLORID. Genes highlighted in green boxes were detected in ≥ 50% of root arrays. Genes in blue boxes were detected in < 50 *%* arrays. Genes and compounds not analyzed in ATH1 arrays are in grey.

If we analysed one-by-one the data and focused on master flowering-time genes that are highlighted in flowering snapshots (Bouché et al., 2016), we found that most of them were actually not expressed in the roots or at least did not pass the filter setting of being detected in at least 50% of the available root transcriptomes (Figure 1C). In the photoperiod pathway, *CO* and *FT* were not at all detected in the dataset; only *GIGANTEA (GI)* was, which mediates between the clock and *CO* regulation (Mishra and Panigrahi, 2015). The flowering activation complex component FD and its paralogue FDP were not hit on the analysis either (detected in 5% of the arrays only). In the aging pathway, *MIRNA* genes were not analysed on ATH1 arrays, but their *SPL* targets involved in flowering were not found in the majority of root microarrays. In the vernalization pathway, *FLC* was detected in 11% of the arrays only. As could be expected, flower meristem identity genes *LFY* and *APETALA1 (APl)* were not detected at all but the upstream MADS box gene *SOC1* was expressed in 42% of the array.

The only pathways whose key regulators are expressed in the roots are the sugar pathway, as *TPS1* was detected in 81% of the arrays, and the ambient temperature pathway, with *SVP* and *FLOWERING LOCUS M (FLM)* coming up in 73% and 51% of the arrays analysed, respectively. This finding makes sense since all plant parts undoubtedly sense sugars and surrounding temperature, including the roots.

### Root transcriptome changes during the induction of flowering

To identify new candidate genes expressed in the roots and potentially involved in flowering, we analysed root transcriptome during the induction of flowering (Figure 2). Plants were grown in hydroponics for 7 weeks under 8-h short days (8-h SD) and then induced to flower by a single 22-h long day (22-h LD), as described in Tocquin et al. (2003). We harvested roots 16 and 22 h after the beginning of the 22-h LD and at the same times in control 8-h SD. We chose these timing points to target early signaling events of floral induction. Two weeks after the experiment, we dissected the remaining intact plants to check that those exposed to the 22-h LD had entered floral transition whereas the 8-h SD controls were still vegetative (Figure 2A). Three independent experiments were performed and used for a transcriptome analysis with Arabidopsis ATH1 genome arrays; the raw results had been included in the data mining reported above. A total of 10,508 AGI loci passed filtering criteria (see Material and Methods) and thus were considered as being expressed in the roots in our experimental system. These 10,508 loci included 168 flowering-time genes, among which 152 were common with the subset revealed by the global data mining shown in Figure 1, somehow confirming these results. Sixteen additional flowering-time genes were then expressed in our experimental set-up, and hence may be regulated by plant age or growing conditions (Supplemental Table 3). Among them, we found the floral integrator *SOC1* and two flowering-time genes involved in the control of meristem determinacy: *TERMINAL FLOWER 1 (TFL1),* a gene of the same family as *FT* but repressing floral transition in the shoot apical meristem (Kobayashi et al., 1999) and *XAANTAL2 (XAL2,* also named *AGL14),* a gene involved in shoot and root development (Garay-Arroyo et al., 2013; Pérez-Ruiz et al., 2015).

**Figure 2.**
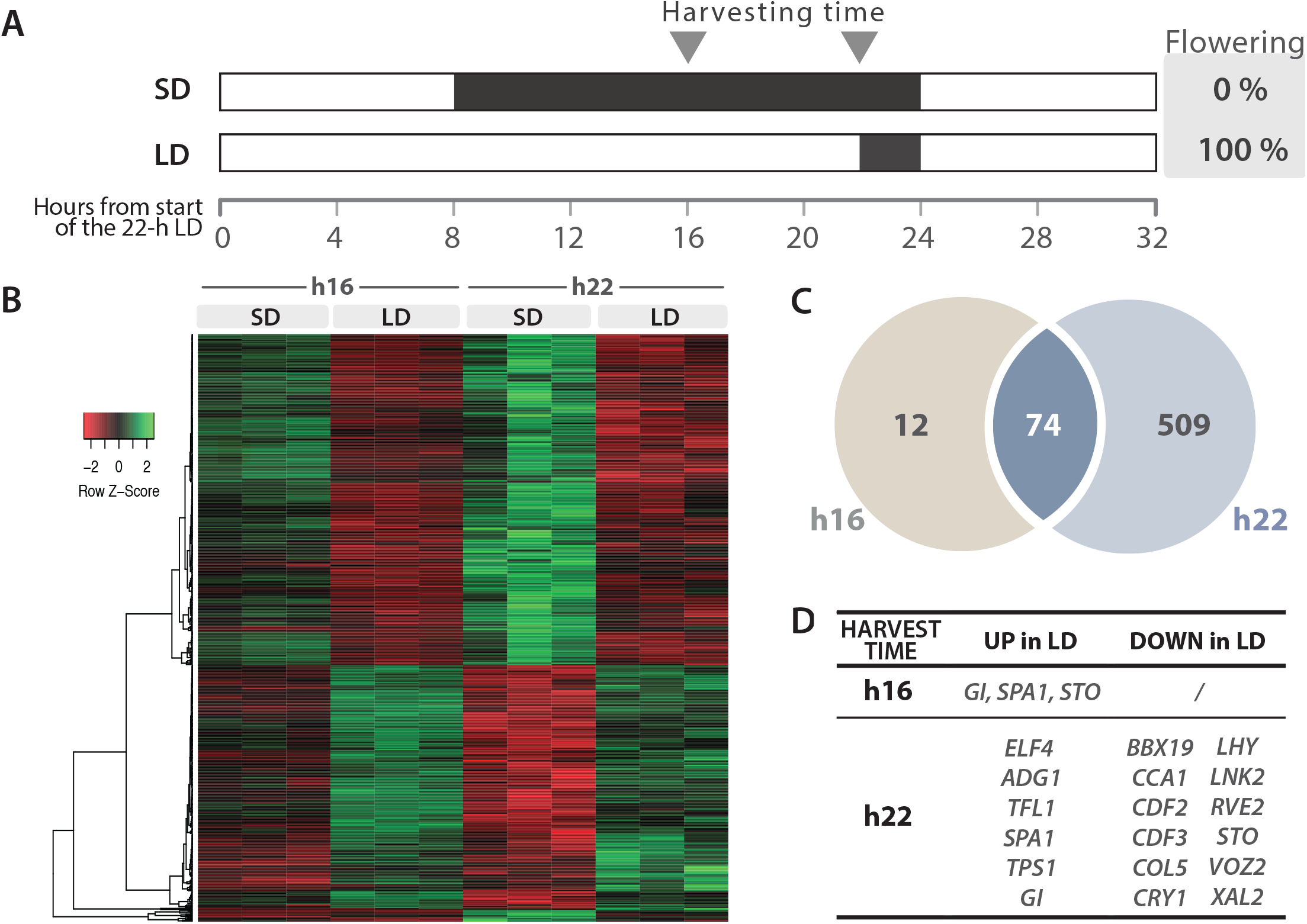
Root transcriptome changes during the induction of flowering by a single 22-h LD. **A**. Experimental design. The proportions of plants having initiated flower buds two weeks after the experiment are shown on the right. **B**. Heatmap of the differentially expressed genes (adjusted p-value ≤ 0.01; fold-change ≥ 2) showing three independent biological replicates per condition. Low expression levels in red, high expression levels in green. Relative expression values are scaled per transcript (lines). **C**. Venn diagram of differentially expressed genes at both sampling time points. **D**. List of differentially expressed flowering-time genes.

The root transcriptome was found to undergo numerous changes during the inductive LD. At h16, *i.e.* 8 hours from the extension of the photoperiod, 86 differentially expressed genes were identified in the roots and at h22, the number had increased to 583 (Figure 2C). We considered genes as being differentially expressed when the adjusted p-value was ≤ 0.01 and fold-change ≥ 2. The heatmap shows that most changes occurring at h16 were actually amplified at h22 (74 of the 86 differentially expressed genes) (Figure 2B) indicating that the experimental design targeted early events. In total, 595 differentially expressed genes were identified in the roots (Supplemental Table 3) among which 18 known flowering time genes allocated to the photoperiod pathway, the circadian clock and the sugar pathway (Figure 2D). This number thus represented about 10% of all the flowering time genes detected in the roots by data mining.

Members of the photoperiodic pathway included negative regulators of CO: *CYCLING DOF FACTOR2* and *3 (CDF2/3), B-BOX DOMAIN PROTEIN 19 (BBX19)* and *SUPPRESSOR OF PHYA-105 1 (SPA1)* but whereas *CDF2/3* and *BBX19* were down-regulated in LD, *SPA1* was upregulated. Two positive regulators of CO were also up-regulated: *GI* and the blue-light photoreceptor gene *CRYPTOCHROME1* (CRY1). Two CO-like genes – *CONSTANS-LIKE5 (COL5)* and *SALT TOLERANCE (STO)* – were down-regulated at h22 in LD, as well as the gene encoding the phytochrome B-interacting protein *VASCULAR PLANT ONE ZINC FINGER PROTEIN 2 (VOZ2).*

Among clock components, several morning genes – *CIRCADIAN CLOCK ASSOCIATED1* (*CCA1*), *LATE ELONGATED HYPOCOTYL (LHY), NIGHT LIGHT-INDUCIBLE AND CLOCK-REGULATED2 (LNK2),* and *REVEILLE2 (RVE2)* – were repressed at h22 in LD. On the opposite, two evening genes were upregulated: *GI* and *EARLY FLOWERING 4 (ELF4).*

The increase in photoperiod also induced the expression of two sugar metabolism-related genes: *TPS1* and *ADP GLUCOSE PYROPHOSPHORYLASE1 (ADG1),* encoding a subunit of ADP-glucose pyrophosphorylase (AGPase). Finally, we found that the expression of two genes involved in the control of meristem fate was also altered: *TFL1* was upregulated in LD whereas *XAL2* was repressed at h22.

### Differentially expressed genes are enriched in phloem tissue

The list of 595 differentially expressed genes was thereafter submitted to different tests to see whether particular networks emerged. We performed three different searches based on (i) tissue enrichment, (ii) gene ontology and (iii) promoter sequences (Figure 3).

**Figure 3.**
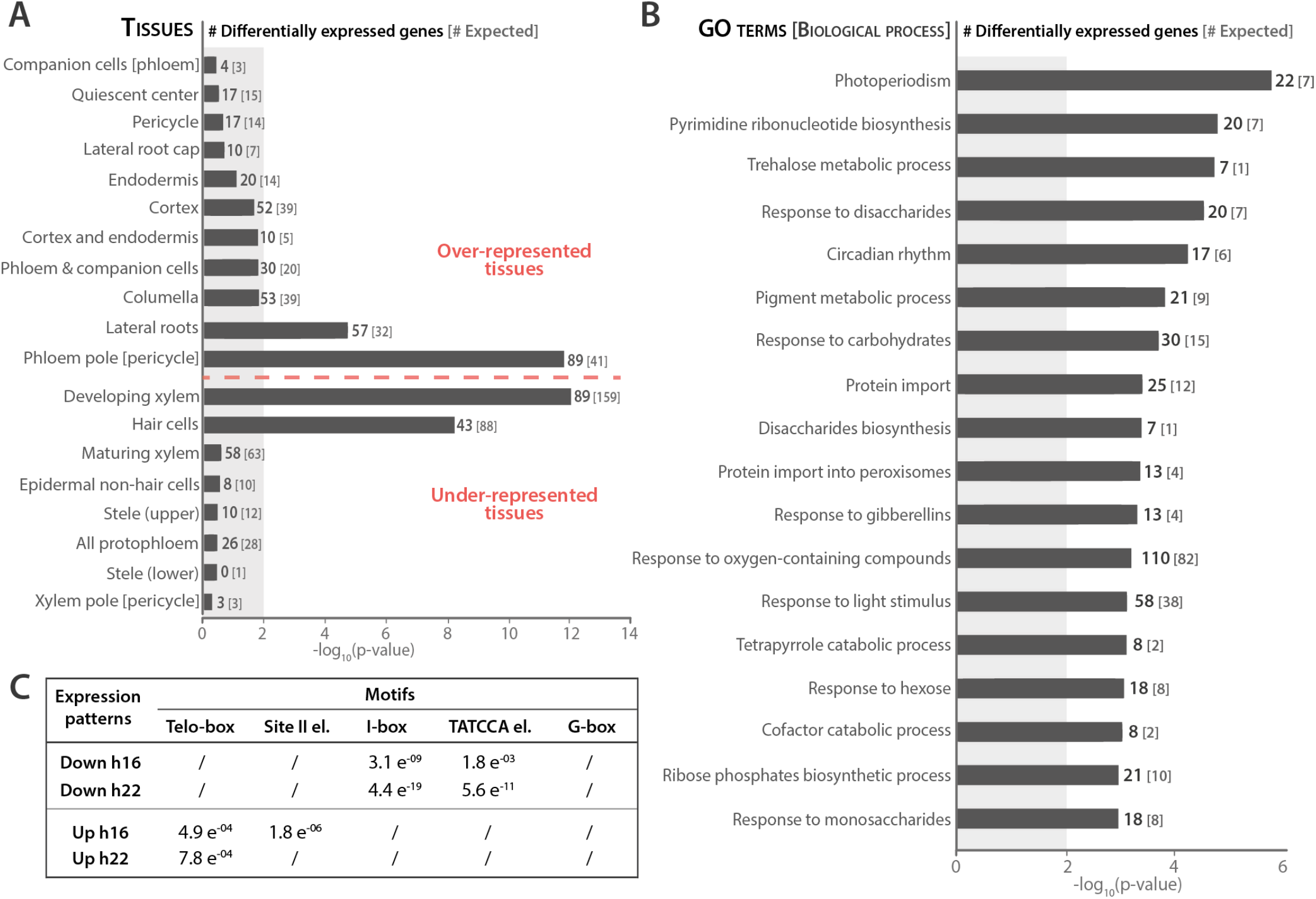
Enrichment analyses of the 595 genes differentially expressed in the roots during an inductive 22-h LD. **A**. Tissue enrichment. For each gene, expression was localized in the tissue where Brady et al. (2007) found highest transcript level. In each tissue, the number of differentially expressed genes is indicated in bold whereas the number of genes that would be expected for this dataset is enclosed within brackets. Shaded area shows p-values > 0.01. Over- and under-represented genes are separated by the horizontal dashed red line. **B**. Gene ontology term enrichment in the list of 595 differentially expressed genes. The number of differentially expressed genes experimentally associated with each term is indicated in bold, whereas the number of genes associated with the GO term that would be expected by chance for this dataset is enclosed within brackets. Bars indicate the −log_10_(p-value) for each term (Fisher’s exact test). **C**. Motif enrichment analysis in the −500 to +50 nt region of the genes that were down- or up-regulated at h16 or at h22 in LD. Numbers are the p-values of motifs that were identified as enriched by AME at p < 0.05 in any of the 4 differentially expressed gene subsets. / indicates non-enriched motif.

First, to know in which tissues the differentially expressed genes were enriched, we crossed their list with the tissue-specific root transcriptome dataset published by Brady et al. (2007). As a reference, we used the whole set of 10,508 genes expressed in the roots in our experimental system. We found that while the genes expressed in the roots are mostly detected in xylem and hair cells, this distribution was notably modified in the subset of differentially expressed genes with phloem and lateral root tissues hosting a significant part of the observed changes (Figure 3A).

Second, we performed a gene ontology enrichment test and found that ‘Photoperiodism’ was the most significantly enriched term in differentially expressed genes (Figure 3B), followed by ‘Pyrimidine ribonucleotide biosynthesis’, ‘Trehalose metabolic processes’, ‘Response to disaccharide’ and ‘Circadian rhythm’.

Third, we searched for enriched cis-elements in the promoters of differentially expressed genes by using the tools of the MEME suite software (Figure 3C). Differentially expressed genes were distributed among four subsets corresponding to the expression patterns illustrated in Figure 2B: up or down in LD, at h16 or h22. A *de novo* motif search was then performed with MEME (motif length between 8 and 15 nucleotides) and DREME (motif length ≤ 8) to find the most represented motifs in the promoters of each of the four gene subsets. Based on Korkuc et al. (Korkuc et al., 2014) study, we scanned the regions spanning −500 to +50 nt from the transcription start site of the genes. Among the resulting motifs, we found several close matches to five known cis-elements: the telo-box (AAACCC[TA]), the site II element (A[AG]GCCCA), the I-Box, the TATCCA element, and the G-box (CACGTG). To determine which of these motifs were specifically associated with the four expression patterns, we tested for the enrichment of each motif in the four subsets of differentially expressed genes with the AME tool. We found that both telo-box and site II elements were significantly enriched in upregulated differentially expressed genes, I-Box and TATCCA were rather associated with repressed differentially expressed genes. The G-box was not significantly enriched in any of the subsets (Figure 3C).

### The change in photoperiod affects the root circadian clock

RT-qPCR analyses were performed on selected differentially expressed genes in order to confirm their differential expression (Figure 4). Since several clock genes appeared on the list, we performed time-course experiments to evaluate in more detail to which extent circadian-regulated processes were affected by the photoperiodic treatment. Roots were therefore harvested every 4 h during the inductive 22-h LD and in control 8-h SD.

**Figure 4.**
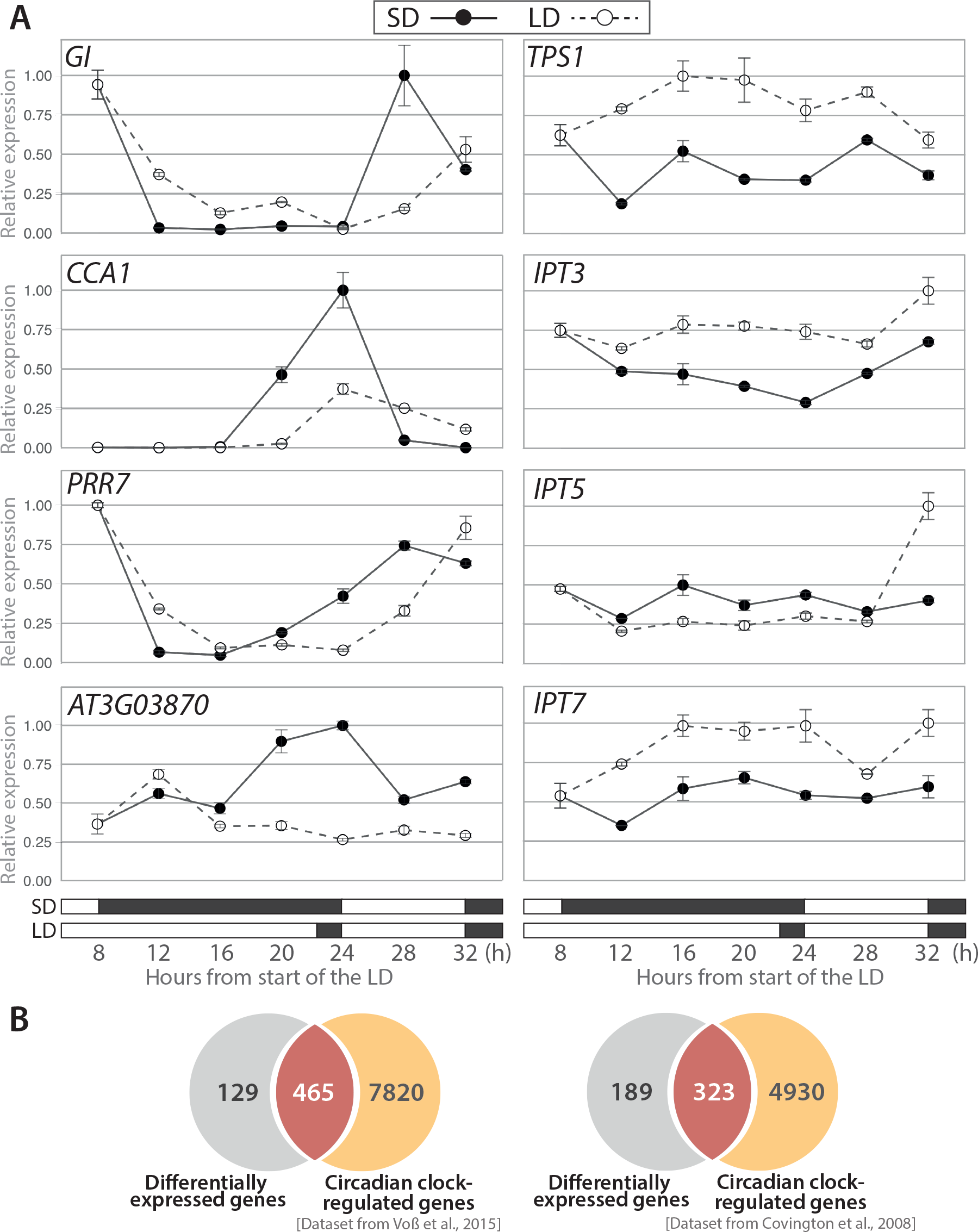
Temporal aspects of transcriptomic changes. **A**. Time-course analyses of candidate gene expression. Relative transcript levels were analysed by RT-qPCR during an 8-h SD (closed symbols) or a single 22-h LD (open symbols). Boxes in the bottom show light (white) and dark (black) periods. Data were normalized using *ACT2* and *UBQ10* genes. Error bars indicate the standard error of the mean for three experimental replicates. Data are from one representative experiment. **B**. Estimate of circadian clock-regulated differentially expressed genes. Venn diagrams showing the overlap between the differentially expressed genes identified in this study and the circadian clock-regulated genes expressed in lateral roots [left; Dataset from Voß et al. (2015)] or in the shoot [right; Dataset from Covington et al. (2008)].

We analysed the expression of *GI, CCA1* and *PSEUDO-RESPONSE REGULATOR 7 (PRR7)* as representative clock genes (Hsu and Harmer, 2014). The 22-h LD caused a 4-h delay in the expression patterns of these three genes, suggesting a phase shift of the circadian clock (Figure 4A, left panel). Since such an effect could globally impact clock outputs, we attempted to evaluate the proportion of clock-regulated genes among the 595 differentially expressed genes. We therefore crossed the list with datasets from transcriptomic analyses of circadian clock-regulated genes in lateral roots (Voß et al., 2015) and shoot (Covington et al., 2008). A large overlap of 78% and 63% was found with these datasets, respectively, revealing that the majority of the differentially expressed genes were indeed regulated by the circadian clock (Figure 4B).

Our analysis also included candidate genes involved in sugar sensing and cytokinin biosynthesis (Figure 4A, right panel). Most interestingly, *TPS1* whose activity is required for flowering in the leaves and in the shoot apical meristem (Wahl et al., 2013) was up-regulated in the roots during the 22-h LD. Our analysis also showed upregulation in LD of two *ISOPENTENYLTRANSFERASE* encoding genes *(IPT3* and *IPT7)* whereas a third one *(IPT5)* did not vary. These results confirmed the microarray data and clearly suggested that sugar signaling and cytokinin biosynthesis were stimulated in the roots in response to the photoperiodic treatment.

### Reverse genetic analysis of differentially expressed genes did not reveal strong phenotypes

We selected a subset of 30 differentially expressed genes for functional analyses, following a number of criteria such as their expression fold change in the microarray analysis, their root-specific expression pattern (inferred from Covington et al. (2008)’s dataset), their putative function or their novelty (Supplemental Table 4). The corresponding mutants available were characterized for two traits: flowering-time and root architecture (Figure 5). Flowering time was quantified as the total number of leaves below the first flower. Surprisingly, only 5 mutants showed an altered flowering time phenotype in LD (Figure 5A). Some of these mutants had been previously characterized such as *gi-2* which, as expected, was very late flowering (Koornneef et al., 1991) and *glycine-rich RNA-binding protein 7* (*grp7,* also called *ccr2*) which was only slightly delayed (Streitner et al., 2008). The cytokinin biosynthesis mutants *ipt3* and *ipt3;5;7* showed an early flowering phenotype but the latter was highly pleiotropic and displayed abnormal growth (Miyawaki et al., 2006). Finally, the mutant for the *AT3G03870* gene of unknown function showed the earliest flowering phenotype, producing 4 fewer leaves than Col-0 WT.

**Figure 5.**
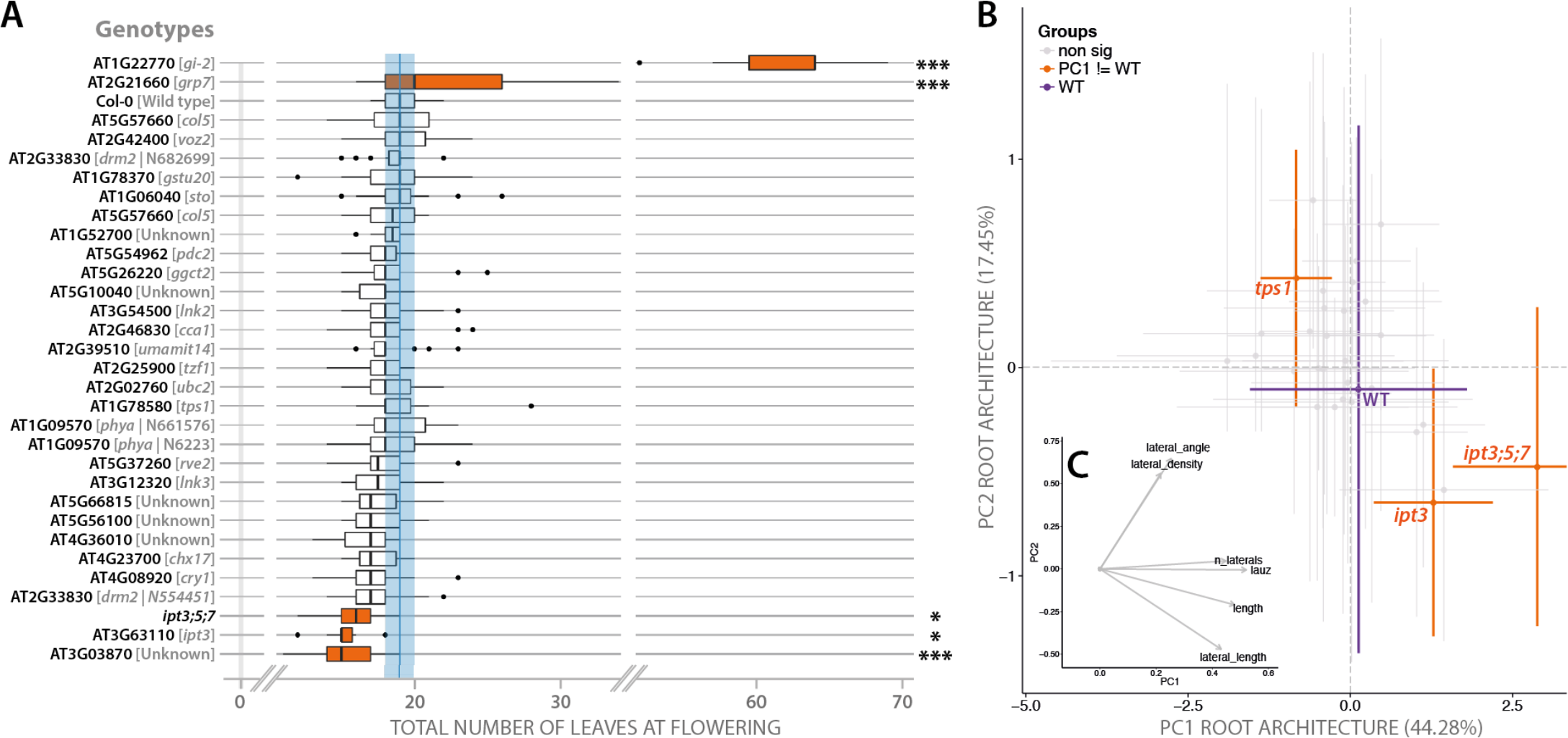
Flowering-time and root architecture phenotypes of selected mutants in 16-h LD. **A**. Total number of leaves below the first flower (n=15). * indicates a significant difference with WT Col-0 (Tukey’s HSD test, p<0.05). *** indicates a highly significant difference with WT (Tukey’s HSD test, p<0.01). WT is shown in blue. **B**. Plot of the first two components of the Principal Component Analysis performed on root system architecture features. **C**. Biplot of the two first components of the PCA. Orange color indicates significant differences with the WT Col-0.

In order to select the genotypes whose root system significantly differed from WT, we performed a Principal Component Analysis (PCA) using the length of the primary root, the length of the apical unbranched zone, the lateral root density, the lateral root number, the total lateral root length and the lateral root angle. The first two Principal Components (PC1 and PC2) were compared using Student tests with a threshold at p<0.01. The selected genotypes were then compared to WT for each variable (t-test, p<0.01) (Figure 5B). The first principal component (PC1), which explained about 45 % of the variability of the dataset, reflects mostly the number of lateral roots, the length of the primary root, the length of the lateral roots, as well as the length of the apical unbranched zone of the primary root (Figure 5C). The PC2 mainly reveals lateral root-related changes, such as their length, their insertion angle on the primary root as well as their density. The *tps1* mutant was affected in PC1 only, showing reduced length of the apical unbranched zone as well as shorter primary and lateral roots. The pleiotropic *ipt3;5;7* triple mutant showed a statistically different PC1, displaying an increased number and density of lateral roots (Chang et al., 2013). The *ipt3* single mutant also displayed a different PC1, albeit with a weaker lateral-root phenotype.

## DISCUSSION

Molecular data concerning the involvement of the roots in the process of flowering are lacking. In this study, transcriptome analyses showed that about 200 genes whose mutation had been shown to alter flowering time are expressed in the roots: 183 were identified in public resources as being expressed in more than 50% of the arrays and 16 additional genes popped-up in our experimental design aiming at analysing root transcriptome at floral transition. This data-crossing relies on an hand-curated database of flowering-time genes that we established recently (Bouché et al., 2016).

The small discrepancy in flowering-time gene numbers found in the two analyses is informative on the fact that some of these genes might be developmentally regulated in the roots. Indeed, most arrays deposited in databases were obtained from a few-day old seedlings whereas we studied mature 7-week old plants. Among the 16 genes expressed in hydroponics but not reaching the 50% threshold in the data mining survey, we found genes regulating meristem determinacy in the shoot: *XAL2* and *TFL1.* Most interestingly, XAL2 is a direct regulator of *TFL1* expression in the shoot apical meristem but both genes have opposite effects on flowering time (Shannon and Meeks-Wagner, 1991; Pérez-Ruiz et al., 2015). Both genes also have opposite effects on root growth: *XAL2* is necessary for normal patterning of root meristem, at least partly through auxin transport (Garay-Arroyo et al., 2013), whereas *TFL1* was recently identified as a repressor of root growth (Lachowiec et al., 2015). We observed that the two genes were differentially expressed in the roots during the 22-h LD, but again in opposite ways: *XAL2* was down-regulated and *TFL1* was up-regulated, a situation that in the shoot would delay flowering and in the root would repress growth. The upregulation of *TFL1* in the root is thus intriguingly similar to what is observed in the shoot meristem where activation of *TFL1* at floral transition is important to counterbalance incoming flowering signals (Jaeger et al., 2013) but whether this is relevant in the root requires further investigation.

In both the global and experimental microarray analyses, the photoperiodic pathway was found to be enriched in the roots and several regulators of *CO* were differentially expressed during the induction of flowering by one LD. Among them we found CDFs and SPA1, involved in the proteolysis of the CO protein. These results are striking since *CO* itself was not detected in the roots, confirming the very low level reported in other microarray studies (e.g. Birnbaum et al. (2003)). In Takada and Goto (2003), several CO::GUS reporter lines showed an expression in the roots while others did not, suggesting that the genomic region in which the transgene is inserted may alter the function of *CO* promoter, which would therefore not reflect its actual expression pattern in roots. Regulators of *CO* thus have other putative targets in the roots, which remain to be discovered. Interestingly, two CO-like genes *(COL5* and *STO)* were found to be downregulated during the inductive LD but whether they share regulatory mechanisms with *CO* is currently unknown.

Some genes of the photoperiodic pathway that are expressed in the roots encode photoreceptors, such as *PHYTOCHROME A-B-C,* and *CRYPTOCHROME1-2* (Supplemental Table 2). Direct light effects on root growth are well documented and several reports therefore recommend to conduct experiments with roots kept in darkness (Yokawa et al., 2013; 2014; Silva-Navas et al., 2015). However, numbers of studies on root architecture in Arabidopsis are performed *in vitro* and routine protocols consist in growing seedlings in transparent Petri dishes with all parts being illuminated. The majority of the root microarrays used for the data mining were obtained from material harvested in these conditions (1,040 out of 1,673 arrays, Supplemental Table 1). We can then speculate that root illumination introduced a bias in the assembled dataset. By contrast, in our hydroponic device, roots were completely in darkness and hence we can assume that any light effect would be indirect. We tested this hypothesis by crossing our dataset with a transcriptomic analysis of seedling roots grown in the dark and exposed to 1-h red light (Molas et al., 2006). After aligning the filter settings of Molas et al.’s analysis with ours, only 55 genes that were differentially expressed after the 1-h red light treatment were detected in the roots in our hydroponics device. Out of them, 11 were differentially expressed during the 22-h LD, including two flowering-time genes: *STO* and *ELF4* (Supplemental Table 5). It is noteworthy however that if both genes are indeed induced by light and interact with different components of light signaling (Khanna et al., 2003; Indorf et al., 2007), they also exhibit circadian expression pattern (Doyle et al., 2002; Indorf et al., 2007), which is the most likely reason why they were differentially expressed in LD.

We indeed estimated that around 70% of the genes that are differentially expressed in the roots during the 22-h LD are regulated by the circadian clock. This proportion is probably overestimated since it was calculated by crossing our dataset with public databases filtered with low stringency tools to retrieve rhythmic gene expression patterns (see Materials and Methods). The clock mechanism was shown in Arabidopsis to rely on three interlocking feedback loops (Hsu and Harmer, 2014). The morning-phased loop comprises *PPR7* and *PPR9* and is activated by CCA1 and LHY; the evening-loop includes EARLY FLOWERING 3 (ELF3), ELF4 and LUX ARRYTHMO, which act together in an evening complex, and other evening genes including *GI* and *TIMING OF CAB EXPRESSION 1 (TOC1).* The central loop makes the link between the two others since TOC1 activates *CCA1* and *LHY* whereas CCA1 and LHY proteins repress *TOC1* (Harmer et al., 2000; Alabadi et al., 2001).

Interestingly, we found that members of the evening loop – *GI* and *ELF4* – were upregulated whereas morning genes such as *CCA1* and *LHY* were downregulated in 22-h LD as compared to 8-h SD. These differential expression levels were recorded at the two time points (h16 and h22) and were probably due to a delay in the expression patterns of these circadian genes upon extension of the photoperiod, as indicated by the time-course analyses (Figure 5) and also reported in other studies (de Montaigu et al., 2015). Such changes might reflect the fact that the circadian clock in plants is entrained to light:dark cycles by photosynthetic inputs. It is known indeed that sugars derived from photosynthesis entrain the circadian clock through morning genes in the shoot (Haydon et al., 2013) and that a shoot-derived photosynthesis product is necessary for the oscillation of the evening genes in the roots (James et al., 2008). Moreover, the circadian clock orchestrates the coordinate adjustment of carbon partitioning and growth rate that occurs in response to photoperiod (Yazdanbakhsh et al., 2011). Consistently, we observed the differential expression of *ADG1,* encoding a subunit of AGPase involved in starch synthesis, and of *TPS1* that catalyses formation of T6P, during the 22-h LD. T6P was found to mediate the sugar-dependent post-translational activation of AGPase (Geigenberger, 2011) and hence upregulation of *ADG1* and of *TPS1* might cooperatively stimulate starch synthesis in the roots during the extension of the photoperiod. Moreover, T6P was found to be positively correlated with rosette growth rate (Sulpice et al., 2014) and to be required in the leaves and the shoot apical meristem at flowering (Wahl et al., 2013). All together, our results suggest that roots are integrated in systemic signaling whereby carbon supply coordinates growth at the whole plant level during the induction of flowering. This coordination possibly involves sugar input to the circadian clock and T6P pathway.

This inference is further supported by our *de novo* analysis of the promoters of genes upregulated at h16 and h22 during the 22-h LD. Both time points revealed an enrichment of the telo-box motif, which is present in the promoter of genes expressed in dividing cells of root meristems and is known to mediate the upregulation of glucose-responsive genes (Rook et al., 2006). The telo-box, which would be part of a midnight regulatory module (Michael et al., 2008), is frequently found associated with other motifs, such as the site II element (Trémousaygue et al., 2003; Zografidis et al., 2014) that we also found in our analysis. The functional relevance of the association between these elements has been demonstrated for the SKIP-mediated control of root elongation (Zhang et al., 2012). Conversely, the promoters of genes downregulated during the 22-h LD were found to be enriched in both I-boxes, which are known to be part of a light regulatory module (López-Ochoa et al., 2007), and in the sugar- and gibberellin-responsive element TATCCA, which is bound by MYB factors (Lu et al., 2002). TATCCA element and G-box were also found to be core components of the sugar response sequence (SRS) in the promoter of a sugar starvation–inducible rice α-amylase gene (Amy3, Lu et al. (1998)). These results support a prominent role for sugars in the control of gene expression during the 22-h LD.

Another coincidence is the enrichment of differentially expressed genes in the phloem tissue of the roots, which is the arrival route of sugars transported from the shoot. For example *IPT3* and *IPT7,* two cytokinin-biosynthesis genes expressed in the root vasculature and the endodermis (Hirose et al., 2008), were differentially expressed during the 22-h LD whereas *IPT5,* which is expressed in the root cap, was not. An increased transport of cytokinins from the roots to the aerial part of the plant would establish a feedforward loop promoting flowering since these hormones are known to activate promoters of flowering in the shoot, such as *TSF* in the leaves and *SOCI* in the shoot apical meristem (D’Aloia et al., 2011). These mechanisms provide a molecular basis to the physiological shoot-to-root-to-shoot loop disclosed in the mustard *Sinapis alba* where sucrose arriving from the shoot induces cytokinin export from the roots to stimulate floral transition (Havelange et al., 2000).

## MATERIAL AND METHODS

### Plant growth

All experiments were performed with *Arabidopsis thaliana* Col-0 accession. The *ipt3* single and *ipt3;5;7* triple mutants were provided by Prof. Tatsuo Kakimoto (Osaka University, Japan) and the *gi* mutant was given by Prof. George Coupland (Max Planck Institute for Plant Breeding Research, Köln, Germany). Other mutants were obtained from the Nottingham Arabidopsis Stock Center (http://www.arabidopsis.info). Accession numbers are provided in the Supplemental Table 4. All seeds, including Col-0 WT, were bulked at the same time to reduce variability. Plants were grown in hydroponic device made of black containers and accessories (http://www.araponics.com). Nutrient solution was a mix of commercial stocks (0.5 ml l^-1^ FloraMicro, FloraGro and FloraBloom; http://www.generalhydroponics.com). Light was provided by fluorescent white tubes at 60 μE.m^−2^.s^−1^ PPFD; temperature was 20°C (day/night) and air relative humidity 70%. For transcriptomic analyses in WT plants, flowering was induced by a single 22-h LD after 7 weeks of growth in 8-h SD and the flowering response was scored as the *%* of plants having initiated floral buds two weeks after the LD (Tocquin et al., 2003). For mutant phenotyping, plants were cultivated in 16-h LD and duration of vegetative growth was scored as the total number of leaves below the first flower (rosette + cauline leaves) to estimate flowering time.

### Microarray analysis

Roots of 18 individual plants were harvested 16h and 22h after the beginning of the inductive LD and pooled. Sampling at the same times in 8-h SD happened during the dark period and were performed under dim green light. Roots were stored at −80°C until used. Tissues were ground in liquid nitrogen and RNA was extracted with TRizol according to manufacturer’s instructions (www.lifetechnologies.com). Before processing further with the RNA samples, we assessed RNA integrity with the Experion^tm^ automated electrophoresis system (www.bio-rad.com). All the samples used for microarray analysis had maximum RNA quality indicator (RQI) values of 10. The RNA samples were labeled using 3’ IVT Expressed kit according to the manufacturer’s instructions (Affymetrix, www.affymetrix.com). Three biological replicates obtained from independent experiments were hybridized on ATH1 Genome arrays (Affymetrix). We analyzed raw data using the limma package (Ritchie et al., 2015). Data were GCRMA-normalized and probeset were filtered for an absolute expression level of at least 100 in ≥ 20% of the arrays. We fitted the data to a linear model using the lmfit() function, analyzed the variance with the ebayes() function, and corrected the p-value for multiple testing using Benjamini and Hochberg’s method (Benjamini and Hochberg, 1995). We considered genes as being differentially expressed when the adjusted p-value was ≤ 0.01 and fold-change ≥ 2.

### *In silico* analysis

Data mining – *In silico* transcriptomic analyses were performed on Arabidopsis Affymetrix ATH1 raw data retrieved from the ArrayExpress database (http://www.ebi.ac.uk/arrayexpress/) using the query "roots". The resulting list was manually sorted to remove experiments lacking comprehensive methodological information. Each experiment was manually curated to select only root-specific raw files. The list of experiments included in the survey is available in supplemental material (Supplemental Table 1). The subsequent data analysis was performed using the R programming language (R Core Team). The "simpleaffy" Bioconductor package V.2.44.0 (Huber et al., 2015; Wilson and Miller, 2005) was used to read the raw data and perform the present/absent call on individual arrays using the detection.p.val() function. Genes were considered as being expressed when p-value <0.01. Within each experiment, we computed the proportion of arrays in which expression of the gene of interest could be detected.

Experimental microarray analyses – The analysis of tissue enrichment was performed using the dataset published in Brady et al. (2007). Each gene represented in the ATH1 arrays was associated with the tissue where its expression level was maximal in Brady’s study. The resulting map was used to localize the genes identified in our study and to calculate their distribution among the different tissues of the roots. This exercise was performed on our microarray analysis with the list of all root-expressed genes (expression level of at least 100 in ≥ 20% of the arrays) or the genes differentially expressed during the photoperiodic induction of flowering (adjusted p-value ≤ 0.01). Using the resulting data, we performed a Fisher’s exact test to determine whether tissues were over- or under-represented in the differentially expressed genes list; the tissues in which the number of differentially expressed genes was higher than the expected value was tested for over-representation while tissues in which the number of differentially expressed genes was lower than the the expect number was tested for underrepresentation (p-value ≤ 0.01).

The Gene Ontology Enrichment analysis was performed using the topGO package V2.20.0 (Alexa and Rahnenfuhrer, 2010) with the annotation of the ATH1 array from ath1121501.db package V3.1.4. We performed a Biological Process (BP) enrichment analysis using the classic Fisher’s exact test (p<0.001). Redundant GO terms were removed. The expected numbers of differentially expressed genes were computed based both on the total number of root-expressed genes (see above) and the number of differentially expressed genes in our microarray analysis.

The analysis of circadian clock-regulated genes exploited datasets obtained in studies of the circadian clock in shoots (Covington et al., 2008) and lateral roots (Voß et al., 2015). To identify the shoot circadian clock-regulated genes, Covington and colleagues analyzed different publically available circadian microarray datasets. We used the list containing the highest number of circadian clock-entrained genes. In Voß’s study, the authors identified highly-probable circadian clock-regulated genes in the roots using three different analysis tools. The list we selected was based on the less stringent parameters, as we included the genes predicted to be clock-regulated by at least one of those tools. When we crossed our experimental list of differentially expressed genes with these datasets, we found that some differentially expressed genes were not represented in Covington’s or Voß’s arrays and hence we excluded them for the comparison.

### RT-qPCR analysis

Roots of 18 individual plants were harvested every 4h during the 22-h LD and at the same times in control 8-h SD. Roots were stored at −80°C until used. Tissues were ground in liquid nitrogen and RNA was extracted with TRizol according to manufacturer’s instructions (www.lifetechnologies.com). RNA samples were treated with DNase (0.2 U DNase μg^-1^). We synthesized first-strand cDNA from 1.5 μg RNA using MMLV reverse transcriptase and oligo(dT)15 according to manufacturer’s instructions (http://www.promega.com). Quantitative PCR (qPCR) reactions were performed in triplicates using SYBR-Green I (http://www.eurogentec.com) in 96-well plates with an iCycler IQ5 (http://www.bio-rad.com). We extracted quantification cycle (Cq) values using the instrument software and imported the data in qbase^PLUS^ 2.0 (http://www.biogazelle.com). A GeNorm analysis (Vandesompele et al., 2002) was performed in a preliminary experiment to identify suitable reference genes. We selected *ACTIN2 (ACT2)* and *TUBULIN2 (TUB2)* as reference genes for root kinetic expression analysis (geNorm M value <0.2). The computed geometric mean of their Cq values was used to calculate the normalization factor, as described in (Vandesompele et al., 2002). The list of primers is available in Supplemental Table 6.

### Root phenotyping

Plants grown for root architecture analysis were sown in vitro on 0.5x MS supplemented with 1% sucrose. Square Petri dishes were used and placed vertically, under 100 μE.m^−2^.s^−1^ PPFD, in 16h-LD. Root pictures were taken every three days using a CCD camera (Canon EOS 1100D with a Canon Lense EF 50mm 1:1.8) and analyzed using the ImageJ plugin “SmartRoot” (Lobet et al., 2011). The resulting root tracing were exported and analysed in R (R Core Team). In order to select the genotypes whose root system significantly differed from WT, we performed a Principal Component Analysis (PCA) using the length of the primary root, the length of the apical unbranched zone, the lateral root density, the lateral root number, the total lateral root length and the lateral root angle. The resulting PC’s were compared using Student tests with a threshold at p<0.01. The selected genotypes were then compared to WT for each variable (t-test, p<0.01). Data visualization was performed using ggplot2 package (Wickham, 2009).

### Cis-elements analysis

For each subsets of similarly controlled genes, we prepared a fasta formated file containing the promoter sequences (-500, +50) obtained from the TAIR10 ftp repository (Lamesch et al., 2012). The analyses were performed using the command line version of the MEME-Suite (Bailey et al. (2015), http://meme-suite.org, version 4.10.0). The parameters for MEME were set as default values, except for: maximum width of each motif: 15 bp; maximum number of motifs to find: 10; background sequences: all TAIR10 promoters (-500, +50). The parameters for DREME and AME were set as default values, with the background sequences being the promoters of the 10,508 genes found expressed in roots in this study.

### Accession numbers

Microarray data are available in the ArrayExpress database (http://www.ebi.uk/arrayexpress) with the accession numbers E-MTAB-4129 and E-MTAB-4130.

## Supplemental data

**Supplemental Table 1**: List of root micro-arrays used for the data-mining analysis. Supplemental Table 2: Flowering-time genes in roots.

**Supplemental Table 3**: List of genes differentially expressed in the roots during a 22h LD. Supplemental Table 4: List of mutants characterized in the present work.

**Supplemental Table 5**: List of genes differentially expressed in the roots during a 22h LD and in Molas et al. (2006).

**Supplemental Table 6**: List of primers used for the RT-qPCR analysis

## ACKNOWLEDGMENTS

The authors would like to thank Kévin Mistiaen for his participation to the setup of microarray analysis pipeline. FB and GL are grateful to the F.R.S.-FNRS for the award of a Ph.D. fellowship (FC 87200) and a postdoctoral research grant (1.B.237.15F) respectively. This research was funded by the Interuniversity Attraction Poles Programme initiated by the Belgian Science Policy Office, P7/29. We thank Prof. Tatsuo Kakimoto (Osaka University, Japan) and Prof. George Coupland (Max Planck Institute for Plant Breeding Research, Köln, Germany) who kindly provided us with *ipt3* / *ipt3;5;7* and *gi* seeds, respectively.

## AUTHOR CONTRIBUTIONS

FB, MD and CP designed the experiments. FB, MD, and ND performed the experiments. FB did the microarray analysis. PT did the promoter analysis. FB and GL did the data analysis and figures. FB, CP, PT and GL participated to the writing of the manuscript. All co-authors read and approved the final version of the manuscript.

## AUTHORS INFORMATIONS

**Frédéric Bouché**, Madison University

**Maria D’Aloia**, GlaxoSmithKline Biologicals

**Pierre Tocquin**, University of Liège

**Guillaume Lobet**, University of Liège

**Nathalie Detry**, University of Liège

**Claire Périlleux**, University of Liège

